# Deciphering the regulation mechanism in biochemical networks by a systems-biology approach

**DOI:** 10.1101/203901

**Authors:** Bernardo A. Mello, Yuhai Tu

## Abstract

To decipher molecular mechanisms in biological systems from system-level input-output data is challenging especially for complex processes that involve interactions among multiple components. Here, we study regulation of the multi-domain (P1-5) histidine kinase CheA by the MCP chemoreceptors. We develop a network model to describe dynamics of the system treating the receptor complex with CheW and P3P4P5 domains of CheA as a regulated enzyme with two substrates, P1 and ATP. The model enables us to search the hypothesis space systematically for the simplest possible regulation mechanism consistent with the available data. Our analysis reveals a novel dual regulation mechanism wherein besides regulating ATP binding the receptor activity has to regulate one other key reaction, either P1 binding or phosphotransfer between P1 and ATP. Furthermore, our study shows that the receptors only control kinetic rates of the enzyme without changing its equilibrium properties. Predictions are made for future experiments to distinguish the remaining two dual-regulation mechanisms. This systems-biology approach of combining modeling and a large input-output data-set should be applicable for studying other complex biological processes.

---

Living systems need to sense environmental signals and use the information to make decisions on their behaviors. The control of behavior is achieved by a myriad of biochemical reactions ^1^, some of which are directly regulated by the sensed signals ^2^. However, exactly which reactions are controlled and how they are controlled by the signal remain challenging questions, especially for complex reaction networks ^3^.

For example, in bacterial chemotaxis ^4,5^, the first stage in signal transduction is from the external stimulus (chemical signal) to a conformational change in the transmembrane chemoreceptors ^6–8^. The second stage is from the conformation state of the receptors to the phosphorylation of the response regulator CheY ^9,10^, which consequently transmits this information to control the bacterial motor and thus the cell motion ^11^. This second stage of signaling relay is performed by the histidine kinase CheA, which contains five functional domains P1-5 ^12,13^. This mechanism, known as the two-component regulation, is prevalent in microorganisms ^14^.

The phosphotransfer from ATP to CheY is performed through several steps internal to CheA ^15^. The internal steps of the kinase activity can be described as chemical reactions involving ATP, CheY, and three sub-units of CheA: P1, P2, and P3P4P5 ^16,17^. These reactants define the chemical states that are connected by bi-directional (froward and backward) reactions forming a reaction network.

The sub-unit P3P4P5 plays a central role in the reaction network, acting as an enzyme that facilitates the phosphotransfer from ATP to P1. For a single enzymatic reaction with one enzyme [*E*] and one substrate [*S*], the experimentally measured production rates for different receptor states are typically fitted by the Michaelis-Menten equation, 
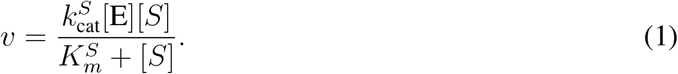

This straightforward analysis can be used to determine how different receptor states regulate the Michaelis-Menten constant 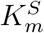 and/or the catalytic rate 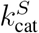. However, such a simple approach fails to reveal the underlying regulatory mechanism if the enzymatic process involves multiple steps with more than one substrate ^18^.

To determine which steps of the reaction network are regulated and how they are regulated is essential to understand the signaling mechanism. However, it is an extremely challenging task given that it is difficult (if not impossible) to probe each individual reaction in the network separately. In this paper, we investigate the regulatory mechanism of CheA kinase activity by modeling the kinetics of the whole enzymatic reaction network under different hypothesis of regulation. The model is fitted to the whole set of experimental data to decide which regulation hypothesis is consistent with the existing data. The model can also provide testable predictions for future experiments to distinguish the remaining hypotheses.

## Methods

Our model is motivated by the recent experimental work by Pan, Dahlquist, and Hazelbauer ^19^ where the authors carried out a comprehensive study of the kinase activity of the functional complex formed by the P3P4P5 units of CheA, the chemoreceptor Tar and the adaptor protein CheW in either nanodisc or vesicle membrane preparations.

The experiments consisted of changing the concentration of one of the two substrates (ATP or P1) while holding constant the concentration of the other substrate for 10 different receptor complex states specified by the receptor methylation level, concentration of aspartate, and membrane preparation (nanodisc or vesicle). From the phosphorylated P1 (P1P) concentration accumulated at time Δ*t*, the average kinase activity 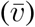 and the average kinase activity per enzyme molecule 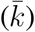 are computed respectively as 
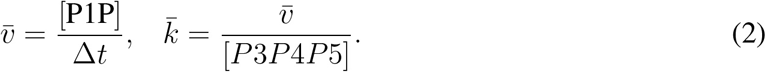

The experimental procedure resulted in 20 “input-output” curves of 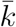 versus [*P*1] or [*ATP*] (two response curves for each of the 10 receptor states). The Michaelis-Menten equation was used in ^19^ to describe each experimental curve separately. Each curve requires one pair of parameters 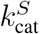and 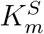, and 40 parameters were needed to fit all the data without any obvious connections among these 40 phenomenological parameters. While CheA kinetics with isolated subunits have already been studied before ^16,17 20^, the recent data from ^19^ are specially suitable for investigating possible receptor regulation mechanisms using our modeling approach because of the large number of receptor states measured in ^19^.

### A network model for CheA enzymatic reaction dynamics

To explain this full set of data within a coherent framework, we developed a simple enzymatic network model to describe dynamics of the enzyme in all possible states in combination with its two substrates/products (ATP/ADP, P1/P1P). The enzyme has two binding sites, one for ATP or ADP and other for P1 or P1P. Each binding site can be in three states: empty, occupied by either the substrate or the product. The combination of these occupancy states results in 3 × 3 = 9 enzyme configurations shown in Fig. 1. The transitions from one state to another form the enzymatic reaction network.

**Fig. 1.**
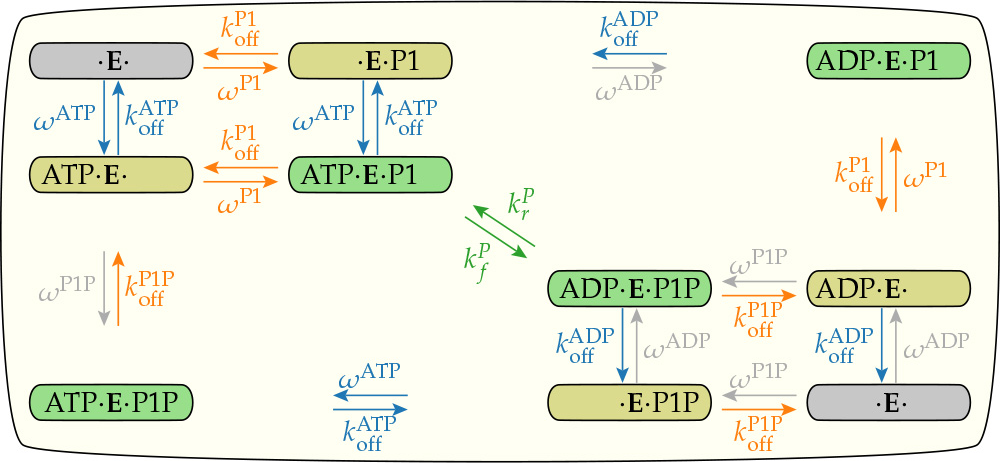
The enzymatic reaction network. The enzyme P3P4P5, denoted by *E*, facilitates the phosphotransfer between its two substrates ATP/ADP and P1/P1P. There are nine states related to the binding of ADP, ATP, P1 or P1P to the enzyme (the empty state is drawn twice). Each pair of vertical or horizontal arrows indicates the binding and unbinding of one substrate to the enzyme. The diagonal arrows in the middle indicate the phosphotransfer reactions. For each substrate *S*(= *ATP*,*P*1), 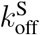 is the dissociation rate and ω^*s*^ is described by Eq. 3. The gray arrows and rates indicate reactions that are negligible because of the low levels of ADP and P1P in the experiments. Reactions that belong to the same regulated mechanism are drawn with the same color, blue for ATP/ADB binding/unbinding, orange for P1/P1P binding/unbinding, and green for phosphorus transfer. The unit of all rates is s^−1^.

The binding/unbinding dynamics of substrate *S*(= *ATP*, *P*1) to the enzyme are described by the dissociation rate, 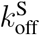 and by the dissociation constant, 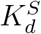, which controls the equilibrium properties of the binding process. For convenience, we define the *on* rate as 
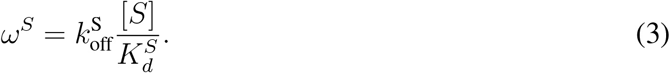

The experimental conditions allow us to assume that [ATP] and [P1] are constant and that [*ADP*] ≈ [*P*1*P*] ≈ 0. This last assumption leads to ω^ADP^ ≈ ω^P1P^ ≈ 0, represented by gray colored arrows in Fig. 1.

The phosphate group transfer rate from ATP to P1 is 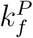. The reverse rate 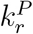 describes the opposite transition. The ratio between these two rates 
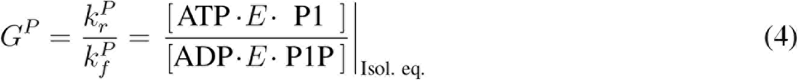
defines the isolated equilibrium between the two states ([**ATP** · **E** · **P1**] and [**ADP** · **E** · **P1P**]). The equilibrium properties depend only on the difference of free energy between the two enzyme configurations, which is given by *k*_*B*_*T* ln *G*^*p*^ with *k*_*B*_*T* the thermal energy unit.

Details of the mathematical formulation of the enzymatic dynamics illustrated in Fig. 1 are given in the Supplementary Material Section S1.

### Incorporating enzyme regulation in the model

Besides defining the chemical reaction network that connects different states of the enzyme, another important ingredient of the model is to specify which reactions (links in the network) are regulated and how they are regulated. Let us characterize the strength of the receptor’s regulatory activity by 0 ≤ *σ* ≤ 1, *σ* = 0,1 correspond to the minimum and the maximum activities respectively.

In our network model of the enzyme kinetics, there are three possible reactions that the receptor activity (*σ*) can affect: the binding/unbinding of ATP/ADP, the binding/unbinding of P1/P1P, and the phosphotransfer between ATP and P1. Regulations in different reactions are labeled by different colors of the reaction arrows in Fig. 1. In this paper, we consider all three possibilities and their combinations to decide what are the minimal sets of regulations needed to explain all the experimental data ^19^.

The exact nature of the regulation determines how *σ* affects the reaction rate *k*_*n*_ for the *n′th* reaction. Here, we consider the simple case where the enzyme has two conformations (active and inactive) and the receptor controls the fraction of time the enzyme spends in each conformation. We further assume the switching between the two enzyme conformations happens at a timescale much faster than that of the other chemical reactions. As a result, the total reaction rates are weighted averages of the “bare” rates in each enzyme state and can be expressed as simple functions of *σ* depending on the nature of the regulation. Specifically, the *σ* dependence is linear if the receptor only affects the kinetic rates without changing the equilibrium properties of the enzyme, but it is a linear rational function if the receptor also changes the equilibrium properties of the enzyme such as the binding energies (see Section S2 in SM for details). We will use our model together with the input-output data to determine the nature of the regulation.

## 1 Results

We search for the regulation mechanism by comparing the best fit of our model to the data under different regulation hypotheses. Specifically, a hypothesis *H*_*i*_ is tested by finding the set of parameters 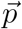 that minimize the mean error χ^2^:

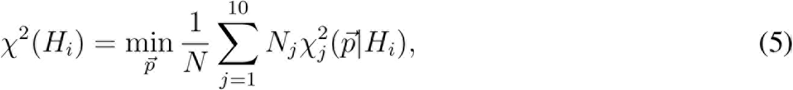

where *j* ∈ [1,10] labels all the 10 individual receptor states characterized by the membrane preparation (“v” for vesicle, “n” for nanodisc), receptor methylation level (EEEE, QEQE, QQQQ), and the ligand (aspartate) concentration (in *μM*). The number of experimental points in each state is *N*_*j*_ (between 12 - 14 data points) and 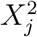 is the mean error of these points, 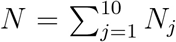 is the total number of data points.

All the model parameters, which are represented by 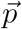, fall into two categories. The first category of parameters include the kinetic rates (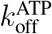, 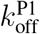, 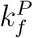) and the equilibrium constants (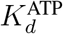, 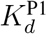, *G*^*P*^), which describe the basic chemical reactions: the binding/unbinding of P1 and ATP to the enzyme and the phosphotransfer between bound P1 and ATP. These “bare” biochemical parameters are the same for all 10 experiments. The second category of parameters are the experiment-specific receptor activity *σ*_*j*_, which can modulate the biochemical rates in experiment *j*(∈ [1,10]). There are a total of 6 + 10 = 16 parameters in our model, which is used to fit all the 20 experimental response curves simultaneously.

Possible regulation mechanisms (hypotheses), represented by *H*_i_, specify how the receptor activity σ affects different reactions in the network. For example, hypothesis *H*_1_ is that the receptor activity only affects P1 binding. For a given hypothesis *H*_*i*_, the total error function is minimized with respect to all parameters and the resulting minimum error χ^2^(H_i_) serves as the error of the hypothesis *H*_*i*_.

This approach enables us to look for the regulation mechanism by searching the hypothesis space systematically. Starting from the simplest regulation rules, we look for significant improvement upon adding new regulatory mechanisms and search for the minimal model(s) that fits all the experimental data. In Figure 2(a), we show the results from some of the tested regulation hypotheses arranged in the legend from the simplest (the top row) to the most complex (the last row). The decomposition of the total error χ^2^ into those from each individual experiments as shown in Fig. 2(a) reveals which experiment(s) invalidates a particular hypothesis and also the possible direction for improvement. We describe our main findings below.

**Figure 2.**
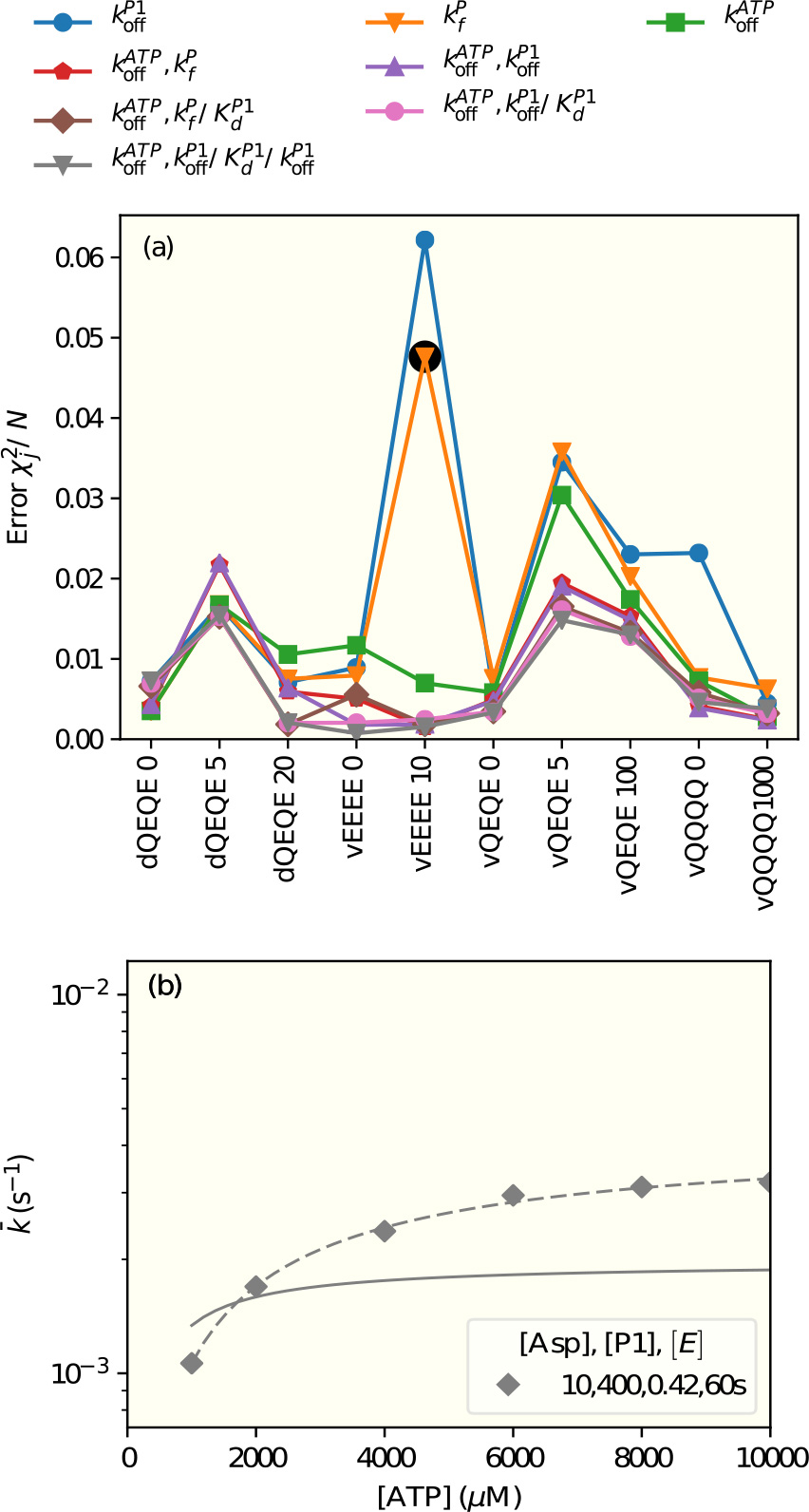
Fitting the model under different regulation hypotheses to the experimental data. (a) The decomposed fitting errors, 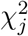, are shown for all 10 experiments listed under the abscissas axis. The different color lines (symbols) correspond to the model with different hypotheses of regulation, which are described in the legend above the plot. The notation of the legend is that the first parameter(s) in the labels are the rates regulated by a. Followed by a slash are parameter(s) that have different values for vesicles and nanodiscs. After the second slash are the regulated parameter(s) with a residual value for *σ* = 0. (b) The actual fit of a failed model to the data for the least active receptor EEEE (highlighted by the back circle in (a)). This failed model under the hypothesis that receptors only regulate the phosphotransfer rate 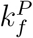 leads to a lower maximum kinase rate and a smaller half-maximum [ATP] concentration in comparison with the experiment (symbols).

### The dual regulation mechanism

As shown in Figure 2(a), none of the three single regulation hypotheses, i.e., regulating the APT or P1 binding or the phosphotransfer rate fits all the experiments. To our surprise, the single regulation of 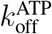 is much better than the other two single regulation mechanisms. The worst performing single regulation hypothesis is regulating P1 binding 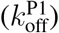, where the errors from several experiments such as VEEEE10, vQEQE5, vQEQE100, and vQQQQ0 are large. For the model with single regulation of the phosphotransfer rate 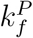, the fitting of vQQQQ0 improves, however the error from vEEEE10 is still large. The detailed reason for this misfit can be understood by comparing the model results with experimental data directly. As shown in Fig. 2(b), the maximum kinase rate and the half-maximum [ATP] concentration for vEEEE10 are both higher in the experiment than in the model. Thus an improvement in fitting this curve seems to suggest an additional regulation of 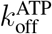.

Based on the results from all the single regulation models, we next tried to combine the different regulations. Quite remarkably, there is a general reduction of errors across most experiments by having 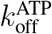 regulation combined with a regulation of either 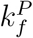 or 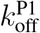. The decomposed fitting errors for these two successful dual regulation mechanisms are shown in Fig. 2(a) (the second row of the legend). However, the dual regulation of 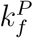 and 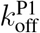 do not improve the fitting (data not shown).

### Receptors regulate the kinetic rates

For a given reversible chemical reaction between two states, the receptor activity (*σ*) can change the energy barrier between the two states and thus change the kinetic rates by the same factor (linearly proportional to *σ*) without changing their ratio, i.e., the equilibrium constants 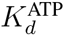, 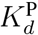 and *G*^*p*^. This is what we used in most of our study. However, we have considered the more general cases when the receptor activity can change the free energy difference between the two states leading to different dissociation constants for the active (*σ* = 1) and inactive (*σ* = 0) receptors and a more complicated (linear rational function) dependence of the forward and backward rates on *σ* (see Section S2 in SM for details). With this new degree of freedom, only slightly improved fittings were achieved as shown in Fig. 2(a) (the fourth row in the legend) for a model with residual activity in P1 binding represented by a second slash followed 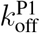 in the legend label. We have also tried allowing this new degree of freedom in the single regulation cases, but the fittings did not seem to improve much if at all (see Fig. S1 in SM).

Our results suggest that receptors mainly regulate the kinetic rates by controlling the energy barrier between two states without changing their free energy difference.

### The difference between nanodisc and vesicle and other observations

In both dual-regulation mechanisms, our model showed that the receptor activities are larger in the vesicles preparation than in the nanodisc by ~ 60 - 90% (see Table 1 and Table S1), probably due to their different receptor cluster structures. We have also considered the possibilities of having different values of 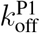, 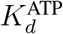, or 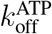 for vesicles and nanodiscs. We found that a modest improvement in fitting can be achieved by having different values of 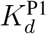 for the nanodisc and the vesicles. These hypotheses are shown in the third row in the legend of Fig. 2(a) by a slash followed by 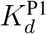. Table 1 and Table S1 show the parameters of one of the two dual-regulation models (regulation of both 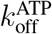 and 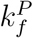) with a higher value of 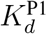 for nanodiscs than that for vesicles by about 50%. The actual fitting of this model to all the experimental data is given in Fig. 3. The parameters for the other possible dual regulation mechanism are given in Table S1 in SM.

**Table 1.** Parameters in the (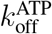, 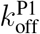) dual regulation mechanism. The fitting isl shown in Fig. 3. The first letter of the label indicates the membrane preparation: nanodisk or vesicle. The following four letters show the receptor’s methylation level.

|  |  |  |  |
| --- | --- | --- | --- |
| $k_{\text{off}}^{\text{P1}}$ | $368 \text{ s}^{-1} \cdot \sigma$ | $K_d^{\text{P1}}$ | $192 / 305(\text{d}) \mu\text{M}$ |
| $k_{\text{off}}^{\text{ATP}}$ | $23 \text{ s}^{-1} \cdot \sigma$ | $K_d^{\text{ATP}}$ | $291 \mu\text{M}$ |
| $k_f^P$ | $7.2 \text{ s}^{-1}$ | $G^P$ | $9.3$ |

Table 1: Parameters in the $(k_{\text{off}}^{\text{ATP}}, k_{\text{off}}^{\text{P1}})$ dual regulation mechanism. The fitting is shown in Fig. 3. The first letter of the label indicates the membrane preparation: nanodisk or vesicle. The following four letters show the receptor's methylation level.
| State | [Asp] | $\sigma$ | State | [Asp] | $\sigma$ |
| --- | --- | --- | --- | --- | --- |
| dQEQE | $20 \mu\text{M}$ | $0.0033$ | vQEQE | $100 \mu\text{M}$ | $0.0059$ |
| | $5 \mu\text{M}$ | $0.066$ | | $5 \mu\text{M}$ | $0.035$ |
| | $0 \mu\text{M}$ | $0.31$ | | $0 \mu\text{M}$ | $0.18$ |
| vEEEE | $10 \mu\text{M}$ | $0.00012$ | vQQQQ | $1000 \mu\text{M}$ | $0.0089$ |
| | $0 \mu\text{M}$ | $0.00081$ | | $0 \mu\text{M}$ | $1$ |

**Figure 3.**
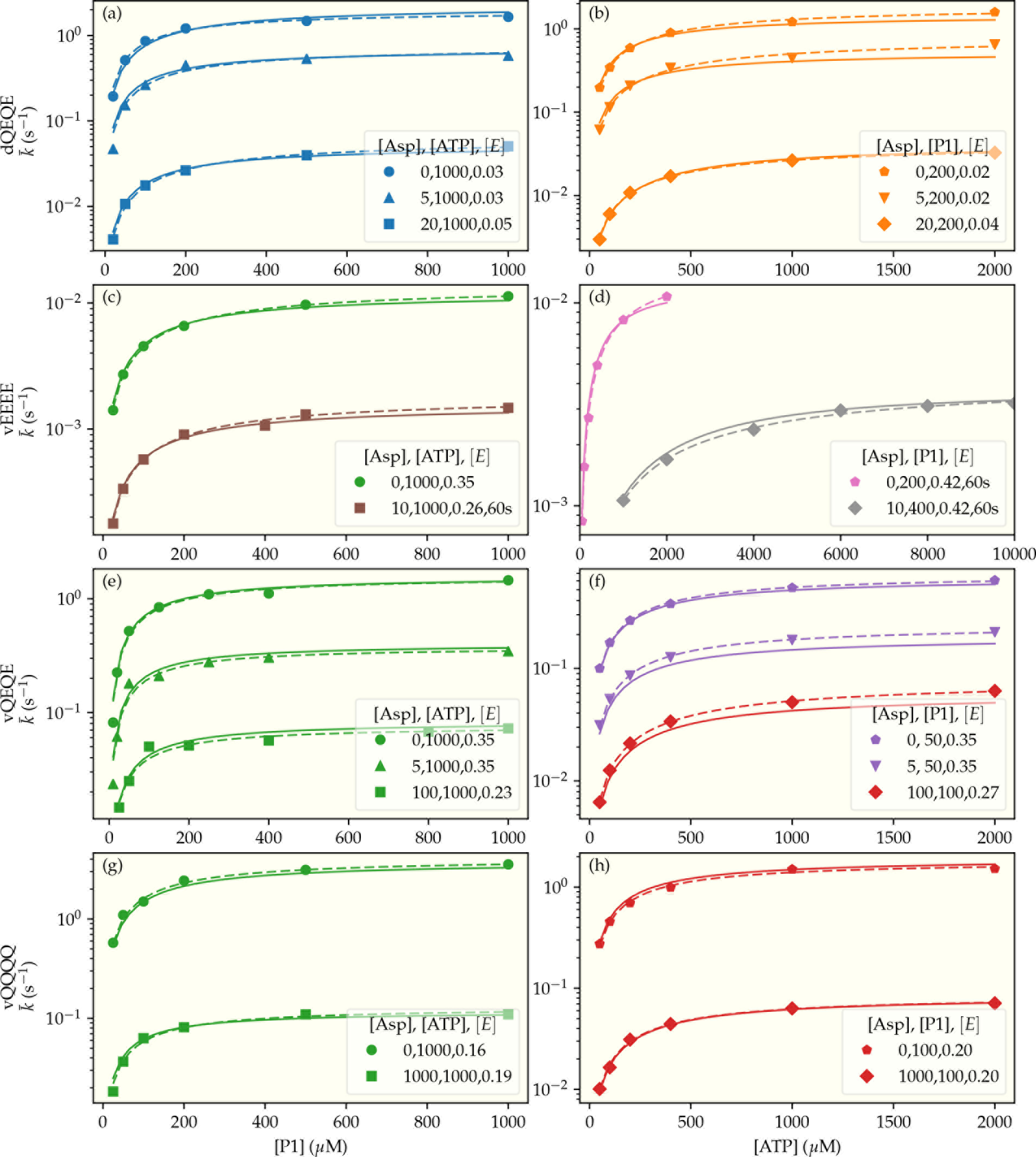
Fitting of our model to all experimental data. The symbols are experimental data of enzymatic phosphorylation rate reported in ^19^. In the left panels, [ATP] is kept constant and [P1] is varied, and vice versa in the right panels. (a)&(b) contain results for receptors in QEQE states in nanodiscs (d). (c-h) contain results from membrane vesicles (v) with the receptors in states EEEE, QEQE, and QQQQ, respectively. The legend in each panel shows the concentrations of aspartate ([Asp]), the constant substrate ([ATP] or [P1]), and of the enzyme P3P4P5 ([E]), in μM. The measurement time was AΔt =15 s for most2aXperiments except for the EEEE receptors when sometimes Δt = 60 s was used as indicated explicitly in the legend. Dashed lines are fitting each experimental curve independently by the Michaelis-Menten equation. Solid lines are from fitting all the data curves together by our model with parameters given in Table 1. The experimental data were provided by the authors of ^19^.

We also explored the possibility of the binding of one substrate depending on the presence of the other substrate as investigated in ^21,22^ and the possibility of 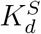 and 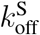 being different for ATP and ADP proposed in ^23^ or for P1 and P1P. However, including these possibilities does not improve the fitting of the available data. Further experiments are needed to explore these more detailed hypotheses.

## 2 Discussion and Summary

In this paper, we developed a simple network model to study the regulatory mechanism of the multi-domain histidine kinase CheA based on recent kinetic measurements. Our best-fit models suggest two possible mechanisms in which the regulatory signal *σ* from the receptor controls the ATP and ADP binding/unbinding rates and either the P1 binding/unbinding rates or the phospho-transfer rates. Previous experimental ^22,23^ and numerical studies have already suggested that the ATP binding was controlled by the receptor activity. However, the dual regulation mechanism discovered is new to the best of our knowledge. Furthermore, our study shows that the receptors modulate the forward and backward rates equally by controlling the barrier between the active and inactive states of the enzyme. The two dual regulation mechanisms are illustrated in Fig. 4. They are consistent with recent molecular dynamics simulations that identified the existence of two states with one of them blocking the access of the substrate to the binding site ^24,25^.

**Figure 4.**
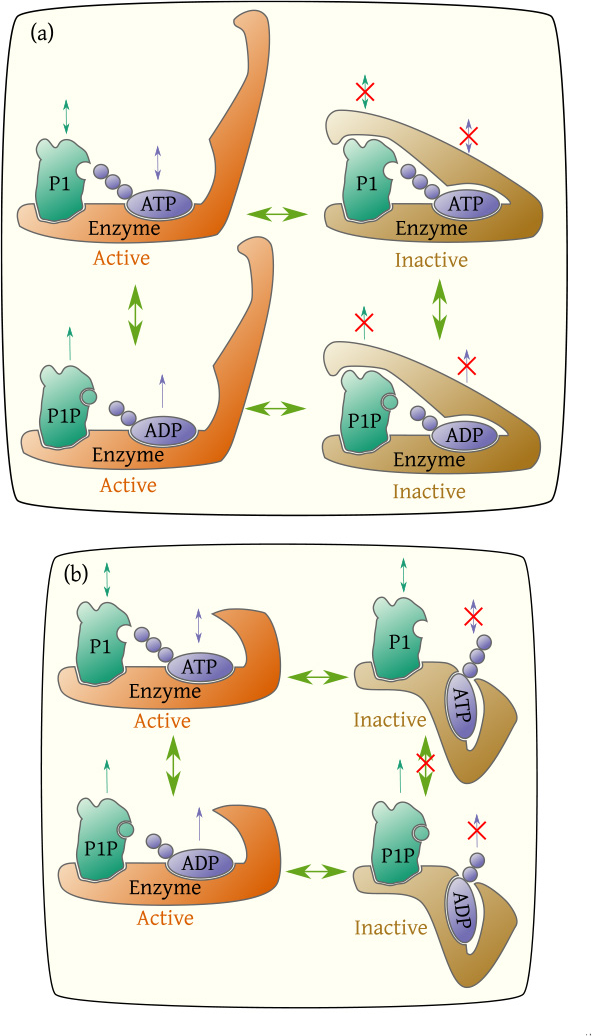
Illustration of the two possible dual regulation mechanisms. The enzyme (P3P4P5) has two states: active and inactive. The receptor activity controls the probability of the enzyme being active. In both mechanisms, the ATP binding site is open in the active state and closed in the inactive state. In addition to regulating ATP binding, the receptor activity also regulates either the P1 binding as shown in (a) or the phosphotransfer between ATP and P1 as shown in (b).

The full network model of the enzymatic reactions can be simplified by exploiting the separation of time scales in the reactions, detailed in the Supplementary Material, sections S3 and S4. These simplifications help us gain more insights on the dynamics of the CheA kinase activity and also lead to predictions for future experiments to further discriminate the remaining regulation hypotheses. We describe some of these insights and predictions below.

### Dynamics of the CheA enzymatic reaction network

Dynamics of the enzymatic network are determined by the transition rates between different states in the network. These different rates give rise to different time scales, which can be regulated by the receptor activity. To demonstrate the importance of the time scales, we plot the phosphorylation rate as a function of time in Fig. 5 for the least active receptor EEEE. We followed the same procedure as in the experiments by premixing P1 with the enzyme and then adding ATP at time *t* = 0. Depending on details of the receptor regulation the phosphorylation rate can either decrease after an initial fast surge (the blue lines) or increase from zero gradually before converging to its steady state value (the orange lines).

**Figure 5.**
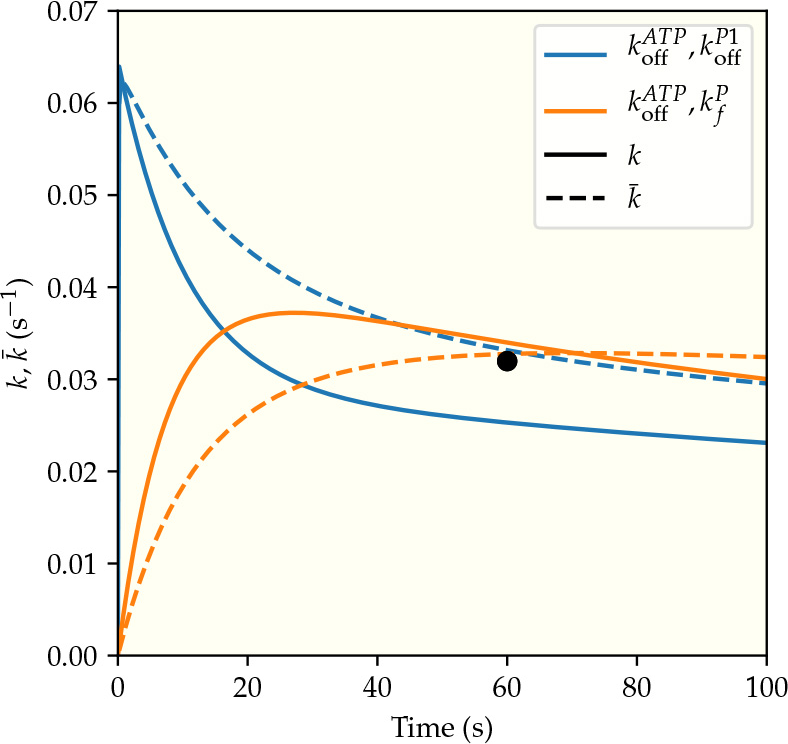
The predicted time dependence of the phosphorylation rates for the two different dual-regulation mechanisms, which are represented by the two different colors (blue for regulating 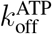 and 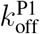 orange for regulating 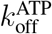 and 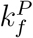). The system studied here contains the least active receptors (EEEE) in membrane vesicles with [Asp]=10 μM, [ATP]=10^4^ μM, and [Pl]=400 ΔM, which corresponds to the conditions of the rightmost point of the lower curve in Fig. 3(d). The solid and the dashed lines correspond to the instantaneous (*k*) and the average 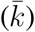 phosphorylation rates respectively. The black circle is the experimental data point from ^19^.

The convergence to the steady state is characterized by the relaxation time, *τ*. Precise calculation of the relaxation time are presented in the SM, but it can be estimated as the inverse of the slowest reaction rate. From the inverse of 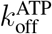 from Table 1, the longest relaxation times are observed for EEEE receptors, estimated as 54 s for [Asp] = 0 and 362 s for [Asp] = 10 *μ*M. This means that the average phosphorylation rates determined in the experiments with Δ*t* = 15*s*, 60*s* are not the steady state rates. We have taken this time-dependent effect into account in all our model fittings explicitly.

Since the Michaelis-Menten (MM) type equation assumes fast equilibration of enzyme binding with its substrates, the MM descriptions of the enzymatic reactions are not correct (at least for the low activity receptors). Nonetheless,the effective (phenomenological) MM fitting parameters may provide useful information regarding the maximum reaction rate and the substrate concentration at which the reaction rate is half of the maximum rate. We determine the values of these effective MM parameters for curves obtained from our model and compare them with those from direct fitting the MM equation with the experimental data. Despite good agreements between the MM parameters from model and data, Fig. 6 also shows the discrepancy of the MM approach. The long relaxation time for small values of 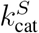 leads to an effective *K*_*m*_ parameter that depends on the measurement time Δ*t*, which is clearly at odds with the MM model assumption.

**Figure 6.**
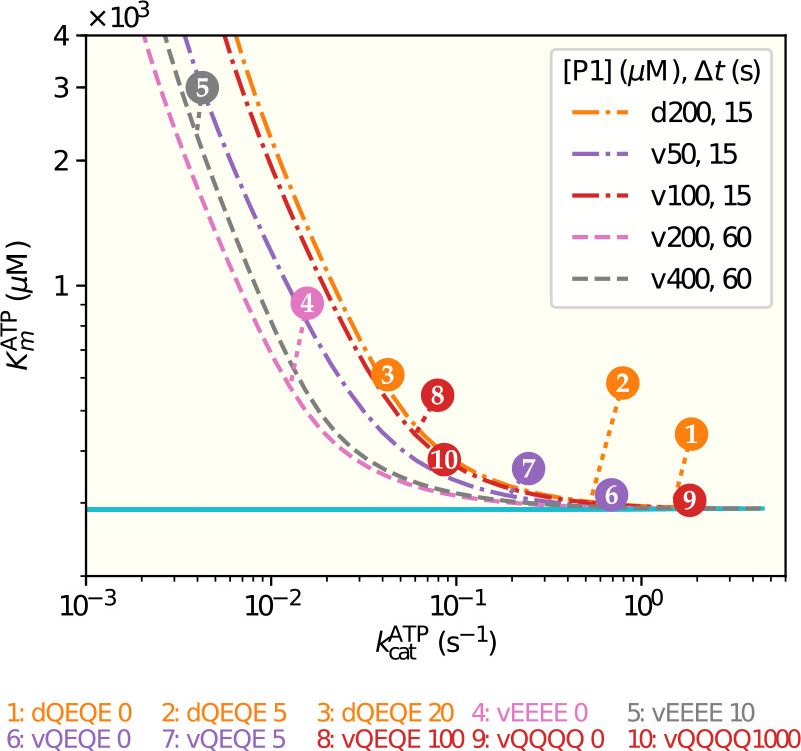
The phenomenological Michaelis-Menten (MM) parameters depend on the receptor activity and measurement time. The pair of the MM parameters (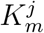,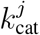) for a given experiment *j* ∈ [1,10] is given by the circle with the experiment number *j*, which is defined the same way as in Fig. 2(a). The dashed lines represent the MM parameter pairs obtained from fitting the corresponding model results with different receptor activities *σ* and at the experimental measurement times (15s or 60s). It is evident that the effective MM parameters depend on the receptor activity and the measurement time Δt· As Δ*t* → ∞ (ploted as a solid line), the effective *K*_m_ approaches a constant independent of *σ* Details on construction of these curves can be found in the Supplementary Material, Section S5.Supplementary Material

### Testable predictions

The rich dynamics of the enzymatic reaction network before it reaches its steady state suggest future experiments that can be used to distinguish the remaining hypotheses. By using our model, we can determine the experimental conditions for which the different hypothesis lead to different dynamical behaviors. The difference in dynamical behaviors is most prominent for the less active receptors (EEEE) where the relaxation times are long.

Fig. 5 illustrates how the two best fitting models, both of which explain the existing experimental data at a particular time Δ*t* = 60*s*, lead to distinctive phosphorylation dynamics due to their different rate limiting steps. In the dual regulation model of ATP and P1 binding, P1 binding is the rate limiting step. As P1 is pre-mixed with the enzyme bypassing this rate limiting step, the phosphorylation rate rises quickly to its maximum before decreasing to its steady state value. In the other dual regulation model of ATP binding and phosphotransfer rates, the combined limitation by ATP binding and phosphorus transfer lead to a much slower initial increase in the phosphorylation rate. Our model also shows that the transient phosphorylation dynamics depend on the initial incubation process (premixing P1 or ATP with the enzyme), see Supplementary Material, Section S6 for details. These predictions of different pre-steady-state phosphorylation dynamics can be tested in future experiments to verify our model and to distinguish the different regulation mechanisms.

In general, to understand the microscopic mechanism in biological systems is challenging given the complexity of the underlying processes and the difficulty in measuring individual reactions. We believe that combining modeling dynamics of the whole reaction network with quantitative system level “input-output” measurements provides a power tool to address this challenge, as demonstrated here in the case of CheA kinase regulation. This systems-biology approach, which includes the development of a mechanistic network model based on key underlying biochemical reactions and searching the hypothesis space by fitting the model results to a large input-output data, should be generally applicable to study other biological regulatory systems.

## Acknowledgements

We acknowledge many fruitful discussions with Drs. Hazelbauer, Dahlquist, and Pan, who provided the raw data from ^19^ shown in Fig. 3. This work is partially supported by a NIH grant (R01-GM081747) to YT.

### Author contributions

YT initiated the project. Both authors contributed equally to the developing of the model, to the analysis of the data and to the writing of the paper. BAM contributed with the developing and running the computational programs.

### Competing Interests

The authors declare that they have no competing financial interests.

## Supplementary Material

### S1 The full mathematical model

The concentrations of the nine enzyme configurations shown in Fig. 1 are described by the vector 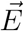, defined as

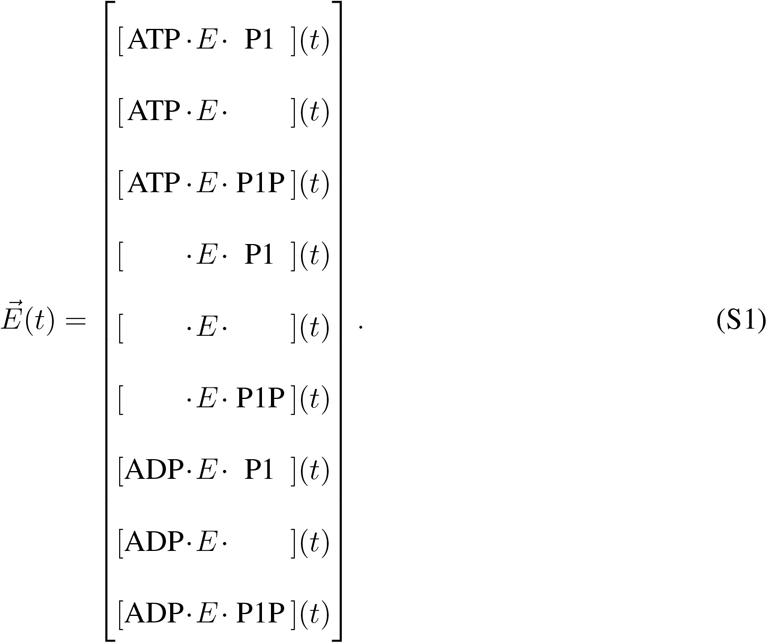

The total enzyme concentration, [*E*], must be equal to the sum of the components of 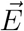, which we define as the norm 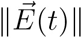. Enzyme conservation imposes

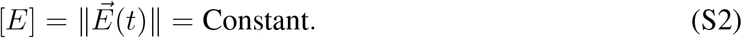

The experimental protocol results in the components of 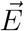 been initially null, except the states [	·*E*·	] and [	·*E*· P1]. We assume these configurations are in equilibrium at *t* = 0, with the values

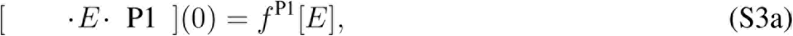

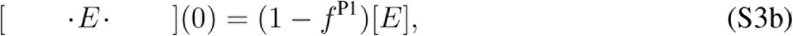

where *f*^P1^ is

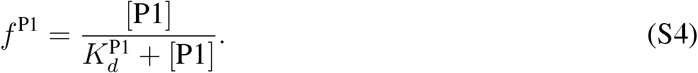

An equivalent approach was used in simulations with pre-mixing with ATP instead of P1.

The evolution of the enzyme vector is described by the equation

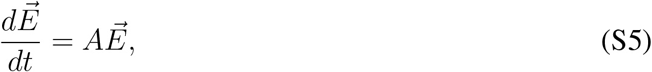

where *A* is the transition matrix

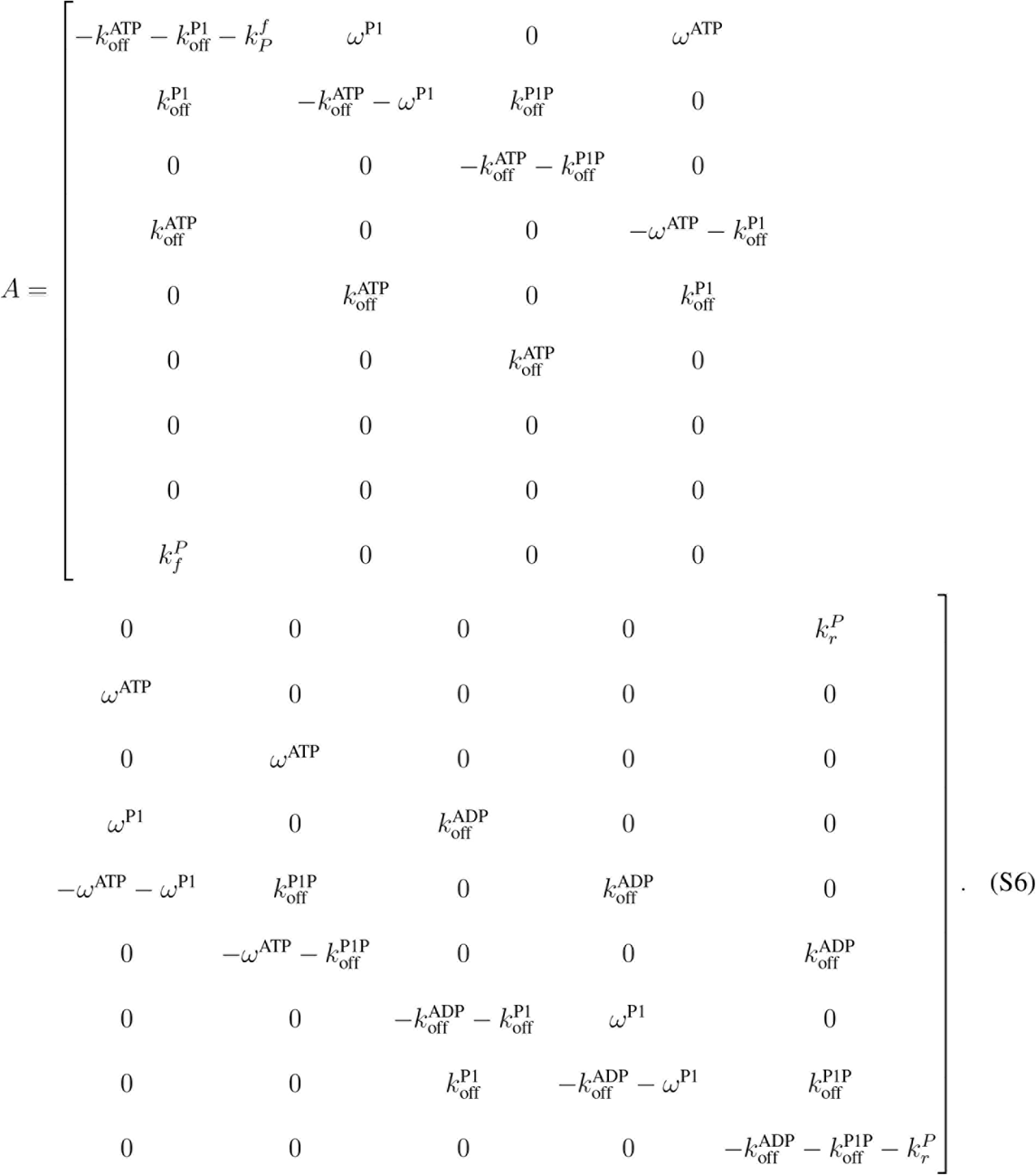

In the general case we should include in the above matrix the rates ω^ADP^ and ω^*P*1*P*^, which were excluded because [ADP] ≈ [*P*1*P*] ≈ 0. Furthermore, the matrix A could be time-dependent, through the rates ω^ATP^ and ω^P1^ which depend on [ATP] and [*P*1]. Due to the minimum consumption of these substances, we assume the matrix to be constant.

The matrix *A* is the infinitesimal generator of the continuous time Markov process describing 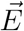 evolution. All eigenvalues λ_*i*_ of such matrices are negative, except for of one which is null, λ_0_ = 0.

By using the eigenvectors 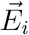 of *A*, the initial state can be written as

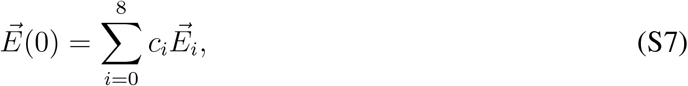

where the coefficients *c*_*i*_ are determined by solving this linear system at time *t* = 0. With these coefficients, the state of the enzyme as function of time can be written as

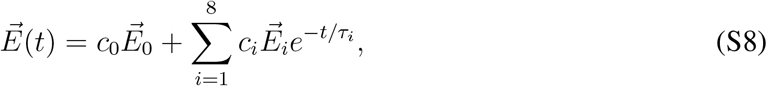

where we used the timescale associated with each eigenvalue,

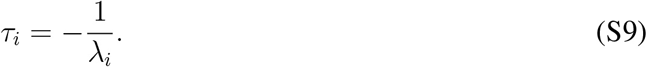

We choose the index of the eigenvalues such that *τ*_1_ > τ_2_ > … > τ_8_.

From Eq. [S8] at *t* → ∞ and Eq. [S2], we conclude that

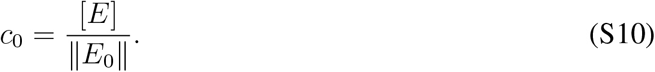

The mean lifetimes of the transients, *τ*_1_, …, *τ*_8_, are associated with the eigenvectors 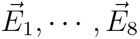. The longest lasting transient is the one with the longest timescale, *τ*_1_, which we will take as the *relaxation time*, τ ≡ τ_1_. It means that the steady state is only reached when *t* ≫ *τ*. The steady state is the eigenvector 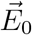 of the null eigenvalue.

The use of the SDS sample buffer in the experiments to stop the chemical reaction means that the measure of [*P*I*P*] include the free and the bound proteins,

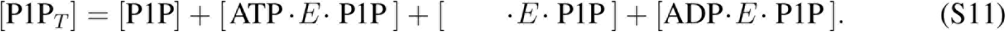

Alternatively, we can calculate [P1P_*T*_] from the net P1 phosphorylation rate

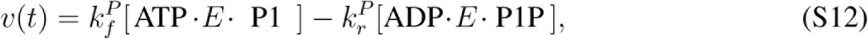

with [ATP ·*E*· P1] and [ADP·*E* · P1P] obtained from Eq. [S8]. We can now calculate the total phosphorylated P1 as

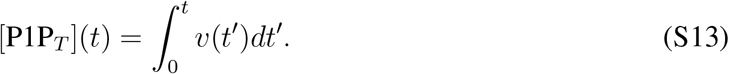

From this, we can write the mean phosphorylation rate as

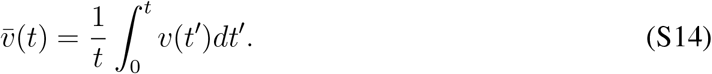

### S2 The models with receptor affecting the equilibrium properties of the enzyme

In the case when receptor activity changes the equilibrium properties of the enzyme such as the binding affinities, substrate binding can be different in the active and inactive state. Therefore, we use a equilibrium model for the substrate binding that involves four states, 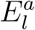, for the substrate bound (*l* = *1*) or not (*l* = 0) and for the active (*a* = 1) and inactive (*a* = 0) states. The free energy of each of these states may be written as

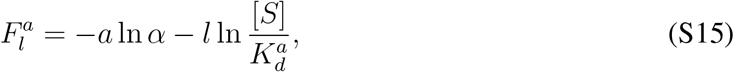

where 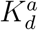 represent the distinct and constant dissociation concentration of the active and inactive states, and *α* is

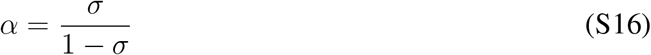

which results in 0 ≤ *σ* ≤ 1 and 0 ≤ *α* ≤ ∈.

The effective binding and unbinding rates for this model is

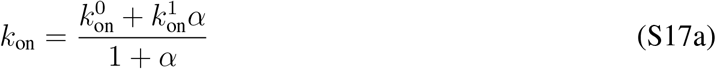

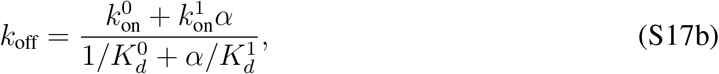

where 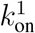 and 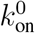 are the binding rate for the active and inactive state, and similarly for the dissociation rate. It is worth mentioning that expressions [S17] require the steady state of the inactive/active switching, not the steady state of the substrate binding. These requirements are satisfied in the experimental conditions.

It is easy to see from Eq. (S17) that when the receptor does not affect the equilibrium binding constant, i.e., 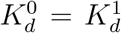, this general model reduces to the case where both the forward and backward rates are proportional to the same factor that is linear in *σ*, which is what we consider in most part of the paper.

Similarly, the free energy of the phosphorus transfer model is

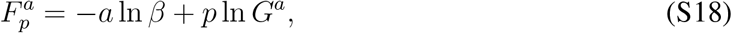

where *p* = 0 is the state with ATP and P1 and *p* = 1 is the state with ADP and P1P, and the constant *G*^*P*^ has now different values in the active and inactive states. The value of *β* is

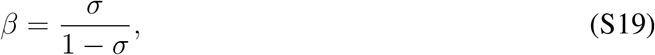

and the effective rates are

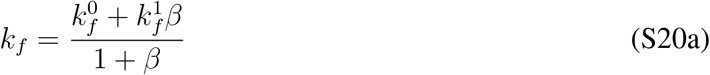

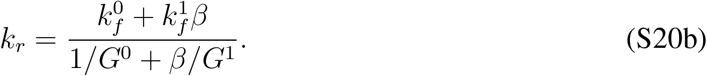

The values of χ^2^ obtained from the fitting with the singly regulated models using the effective rates [S17] and [S20] are shown in Fig. S1.

### S3 Simplifying the model witth 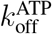 and 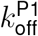 regulation

**The five states model** The full model can be simplified by assuming that states connected by fast reactions and surrounded by slow reactions are in the steady state. The property 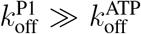 and ω^P1^≫ ω^ATP^ in Table 1 justify assuming the steady state in the binding of P1 to the enzyme.

In the five states model, we group together some pairs of states that differ only by the P1 binding site been empty in one state and occupied by P1 in the other. We use the following convention to represent these grouped states

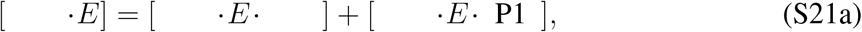

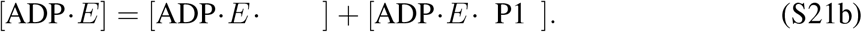

In figure S2(b) we see the states and the reactions of the five states model. The occupancy of the P1 binding site in the grouped states is given by Eq. [S4].

**Figure S1.**
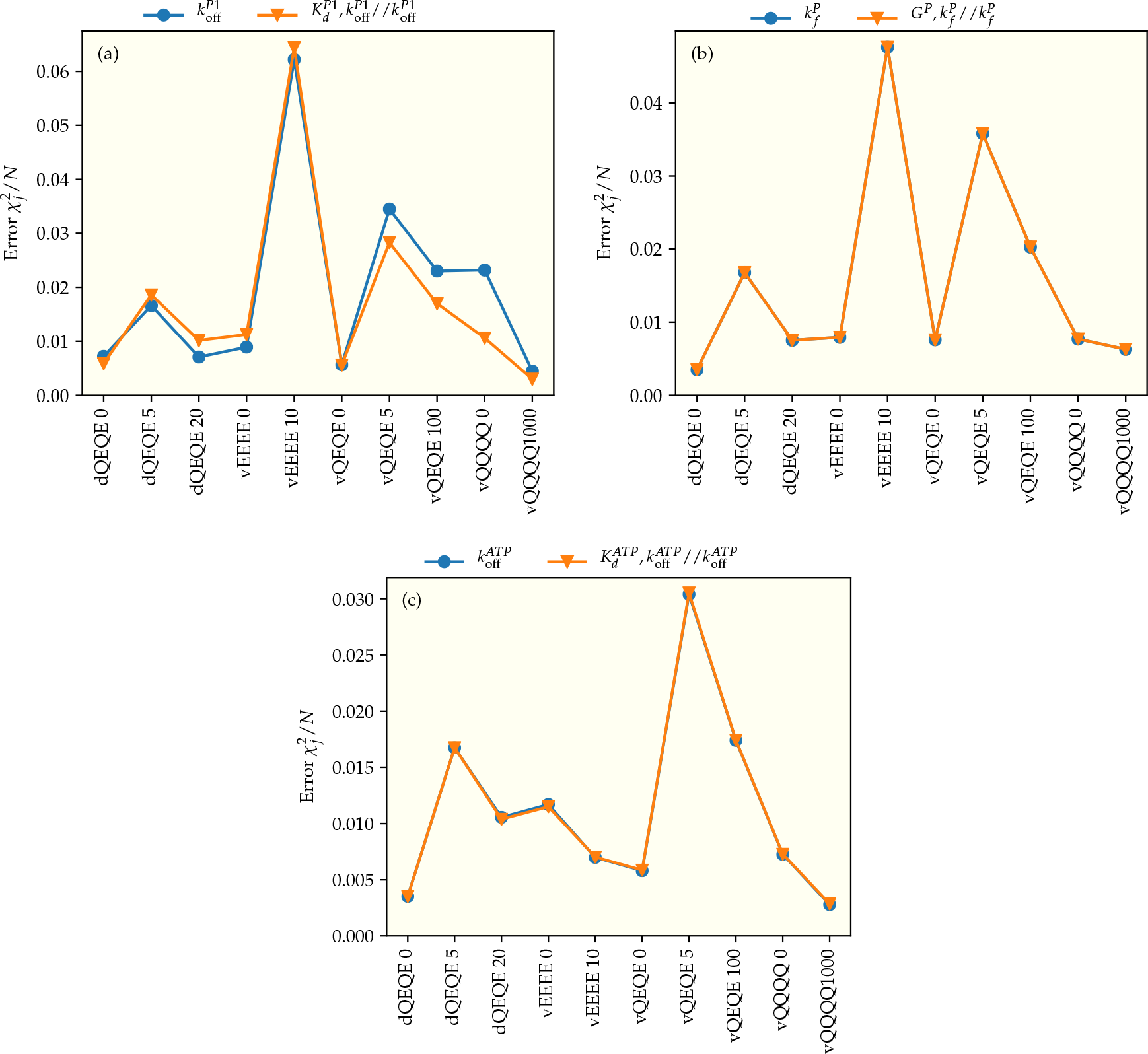
Comparison of models with and without residual activities. Fitting errors in models with (orange) and without (blue) residual activities when the receptor activity regulates (a) the P1 binding, (b) the phosphotransfer rate, (c) the ATP binding rate. The improvements of fitting by including residual activities are minimal.

**Figure S2.**
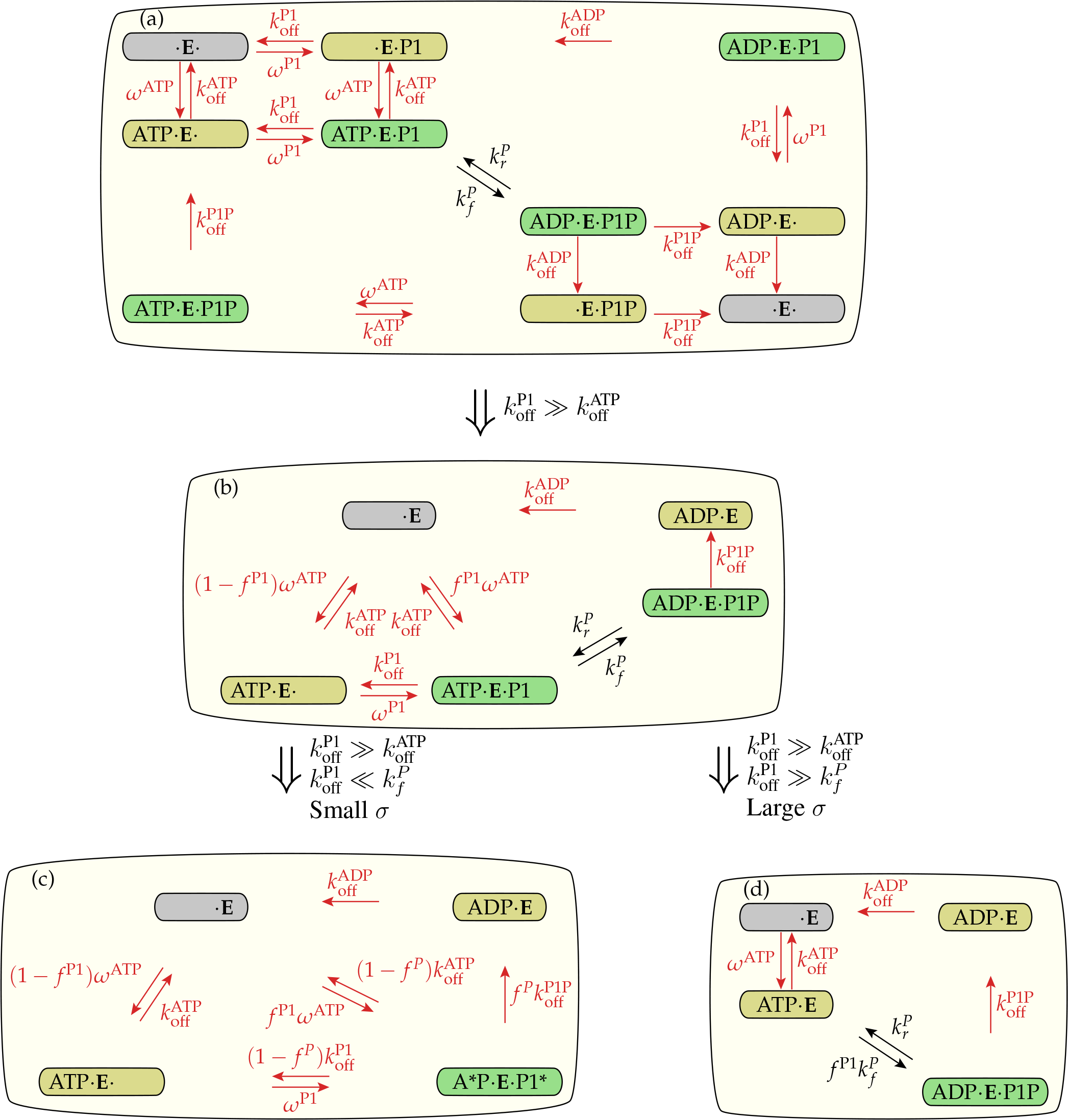
Using parameters from Table 1 to simplify the enzymatic network of Fig. 1. (a) The same as Fig. 1 for the model with regulation of 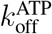 and 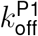, regulation represented by red arrows and rates. (b) The parameters allow disregarding some states and grouping together some states that differ only by the attachment of P1. It results in the five states model, which describe the experimental data as well as the full-model. Four states models can be used when σ is small (c) of large (d).

The much faster unbinding of P1P compared to ADP allow us to disregard the reaction ADP·*E*·P1P ⟶ *E* ·P1P Because of the negligible concentration P1P and ADP the concentration of ATP·*E*·P1P and ·*E*·P1P are also negligible. However we must keep [ADP·*E*·P1P] and [ADP·*E*], which results from the phosphate transfer reaction.

The high ratio 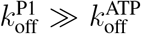 make the five states model a good replacement for the full model in the experiment conditions. It is the starting points for further simplification.

**The complex four states model** If 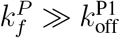 we can define the state

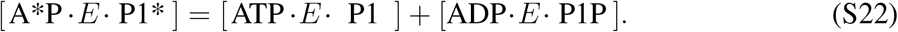

By assuming the steady state of the phosphorus transfer reaction we can write [ADP · *E* · P1P] = *f*^*P*^[A *P · *E*· P1*], with

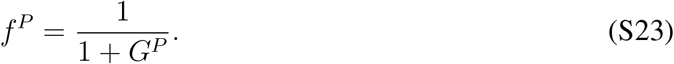

The resulting four states model, shown in Fig. S2(c), can be used when 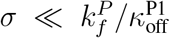 where 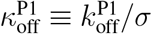

**The simple four states model** If 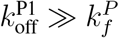 we can assume the stead state of P1 binding to ATP·*E* and define the state

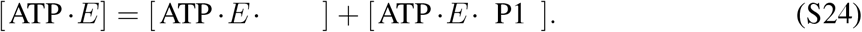

This condition implies in 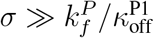

In figure S2(d) we see the states and the reactions of the four states model for large *σ*. The phosphorus transfer reaction is proportional to

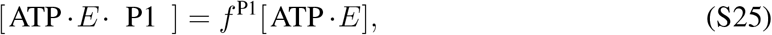

with *f*^P1^ given by Eq. [S4]. Due to the high ratio 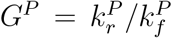 shown in Table 1, the reverse phosphorylation rate is much higher than the forward rate. For this reason we can’t assume instantaneous detaching of P1P and the state [ADP·*E*· P1P] must be explicitly written. The value of 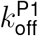 is irrelevant for this model, as long as it is large enough.

The simplicity of the four states model with large *σ* allows obtaining analytic expressions for the Michaelis-Menten parameters, as we do next. The mathematical model of Fig. S2(d) requires a four-component enzyme vector,

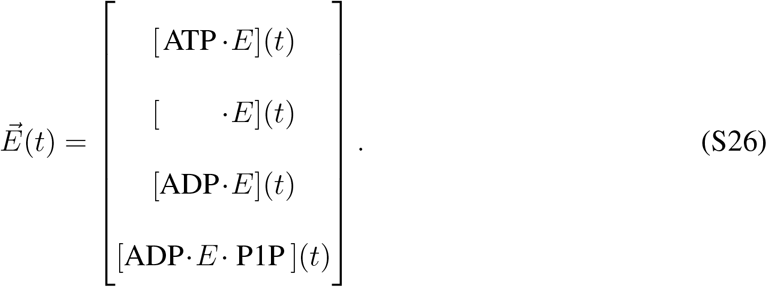

and the transition matrix

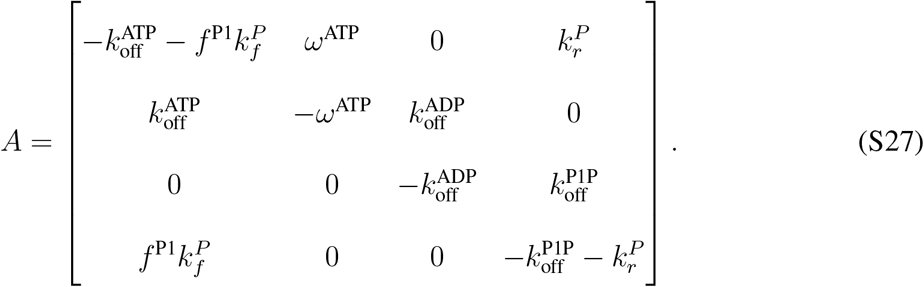

The null eigenvector of this matrix is

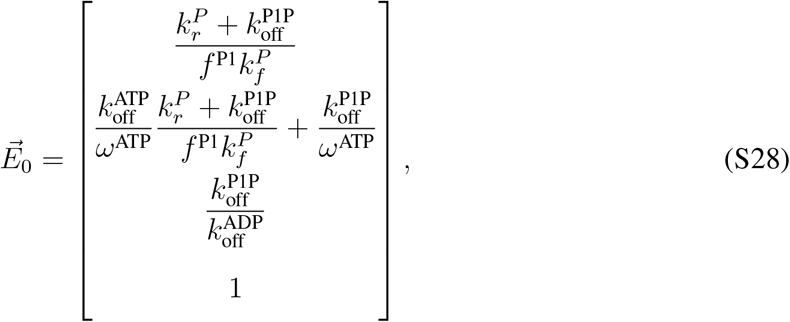

which is also the steady state.

Instead of using Eq. [S12], we can calculate the steady state phosphorylation rate of the four states model as

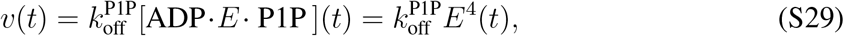

where *E*^4^(*t*) is the fourth component of 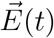 in Eq. [S26]. For the steady state, we can write

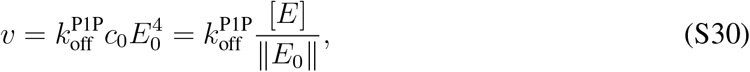

where we used Eq. [S10].

It is straightforward to show that Eq. [S30] can be written as the Michaelis-Menten equation, Eq. [1], either as a function of [P1], with the constants

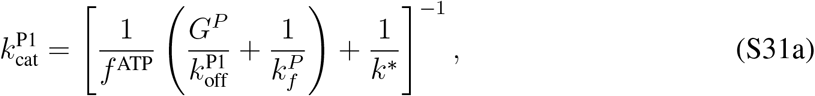

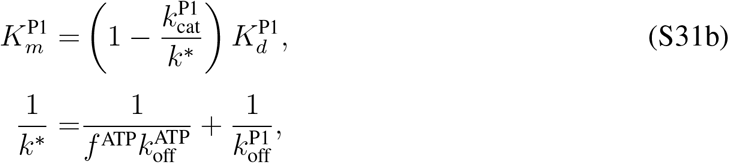

or as a function of [ATP], with the constants

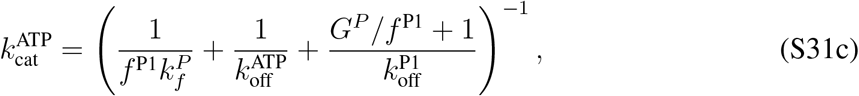

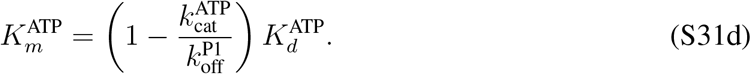

In the above expressions we used the identity 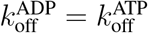 = koff obtained from the fitting.

The concentration of ATP can affect 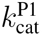 and 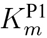 through the parameter *f*^ATP^ and the concentration of P1 can affect 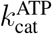 and 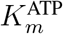 through the parameter *f*^P1^. The four Michaelis-Menten constants can be affected by the regulatory signal a through 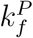 and 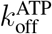.

**Figure S3:**
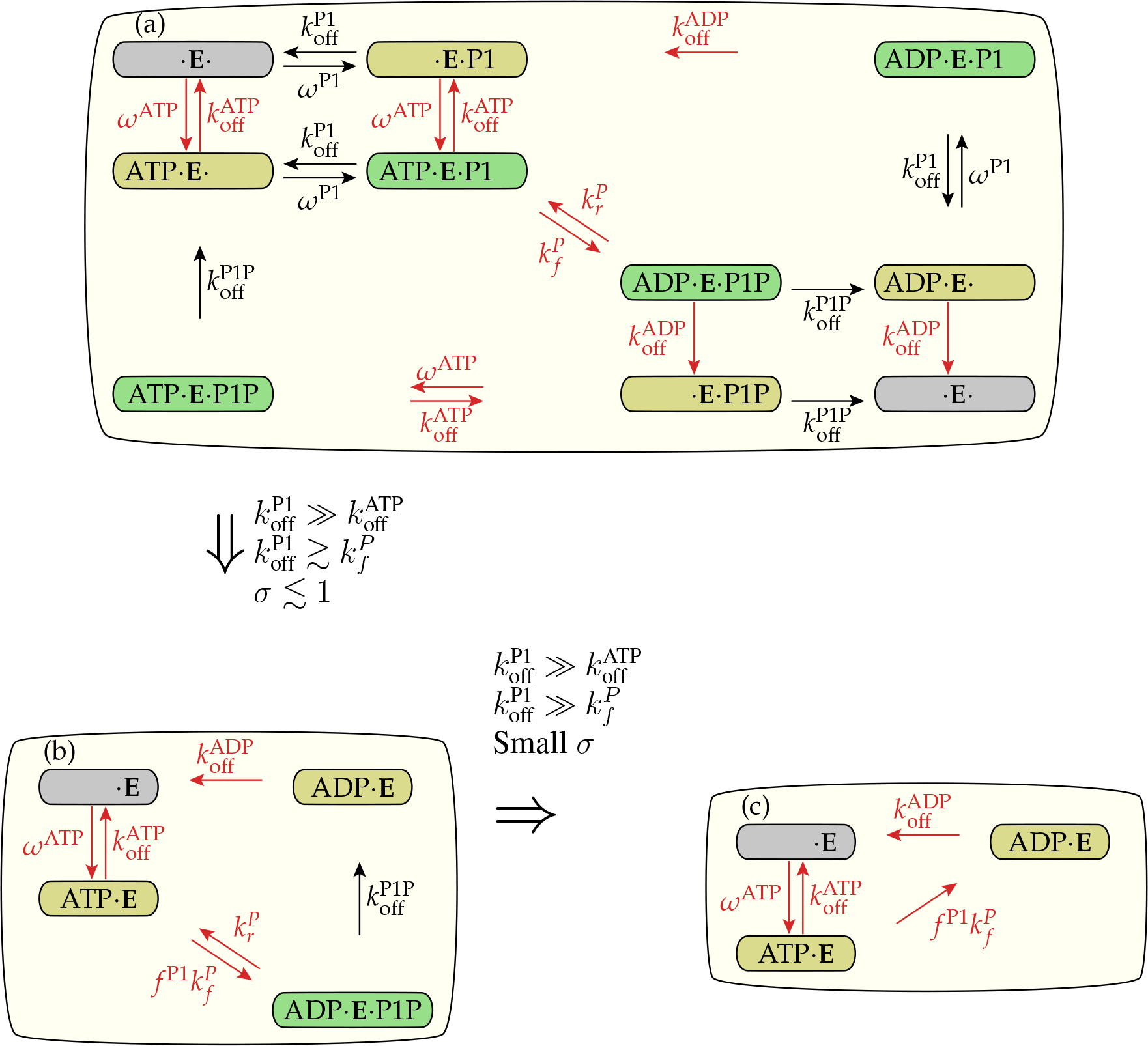
Using parameters from Table S1 to simplify the enzymatic network of Fig. 1. (a) The same as Fig. 1 for the model with regulation of 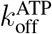 and 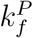, regulation represented by red arrows and rates. (b) By removing the non-essential reactions we can write the core model, which fits the data as well as the full-model. In the core model there are no receptors in the states ATP·E·P1P or ·E·P1P and the small boxes represent the mix of enzymes with empty and occupied P1 binding sites. (c) A simpler model can be used if 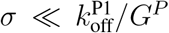. In this case, we can assume that the phosphate is transferred from ATP to P1, but never from P1P to ADP.

### S4 Simplifying the model with 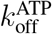 and 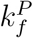 regulation

**The four states model** By observing Table S1 we can see that for this model 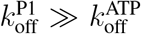 and 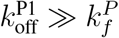, for all values of *σ* < 1. For this reason, the model shown in Fig. S3(a) can be replaced by Fig. S3(b). This model and the model of Fig. S2(d) involve the same chemical reactions and share the same mathematical properties discussed in Section S3-S3

**The three states model** Table S1 shows that if σ is sufficiently small, then we can assume 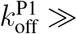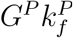. In this case, the four states model of Fig. S3(b) can be further simplified to S3(c). This model requires a three-component enzyme vector,

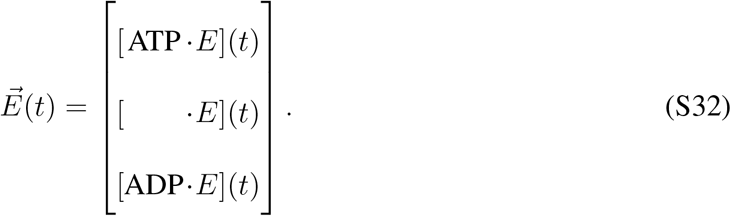

and the transition matrix

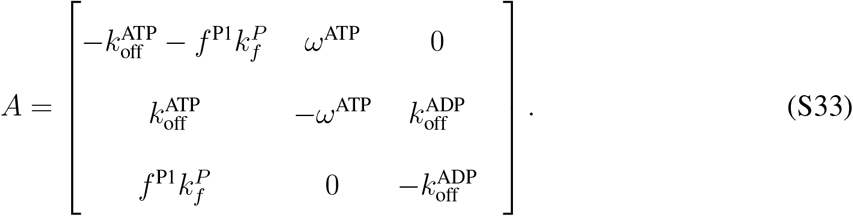

The null eigenvector of this matrix is

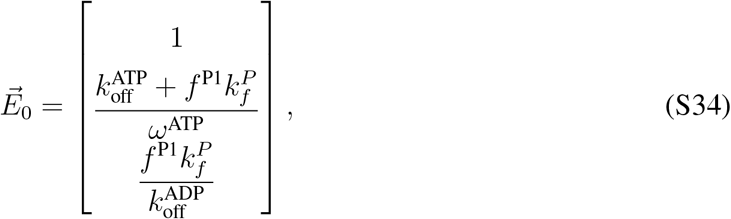

which is also the steady state.

The phosphorylation may be obtained from

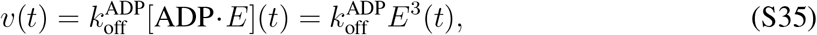

where *E*^3^(*t*) is the third component of 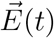 in Eq. [S32]. For the steady state, we can write

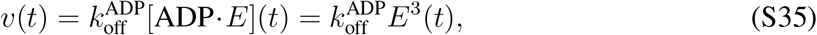

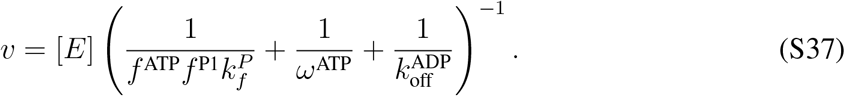

By making 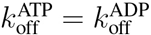 we can write,

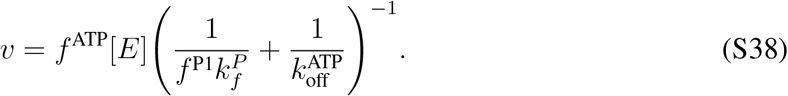

which can also be obtained from Eq. [S30] in the limit 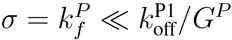. The Michaelis-Menten parameters for this equation are

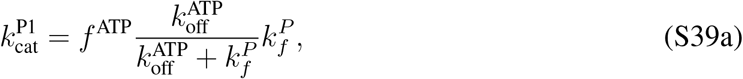

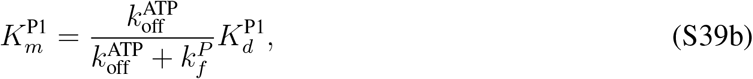

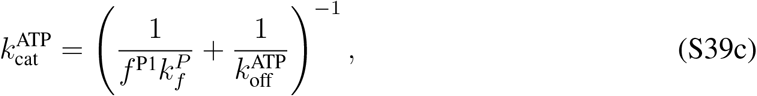

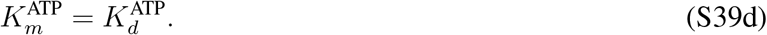

These functions are plotted as the black dotted lines in Fig. 6a-d with the values from Table 1, and perfectly match the exact model for 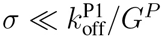.

Due to the simplicity of the minimal model, the non-zero eigenvalues and eigenvectors with finite timescales are easily found when 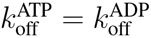,

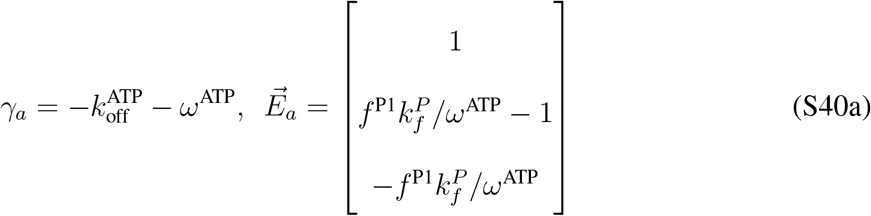

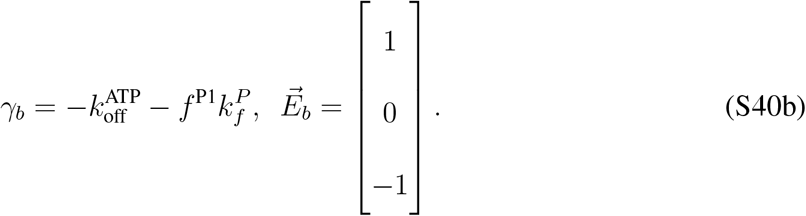

It results in the following timescales

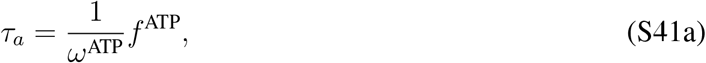

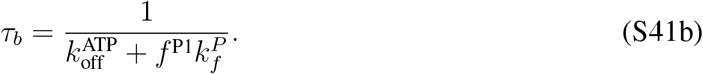

### S5 The effective Michaelis-Menten parameters from simulations of the model

Figure S5 illustrate the dependence of the Michaelis-Menter parameters on the regulatory signal and substrate concentration. The simulations were performed for the same concentrations of the substrate hold constant in the experiments, as indicated in the legends of Fig. 3. Michaelis-Menten equation as a function of the other substrate was used to obtain 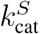 and 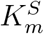.

The values of *σ* are not measured in the experiments. For this reason, it is not possible to compare the experimental results directly with Fig. S5(a-d). To allow this comparison, we combined the points with the same *σ* from the curves of 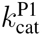 and 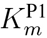 to create the curve of 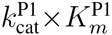 of Fig. S5(e). The same procedure was used to create Fig. S5(f).

### S6 Simulation of the model with premixing of enzyme and ATP

Regulations of 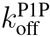 present in one model and absent in the other, can explain the differences caused by incubation (premixing), shown in Fig. S4. The low concentration of P1 and the high concentration of ATP make P1 binding the limiting reaction at the onset of the experiment. For this reason, changes in the regulatory signal will have distinct effects on the phosphorylation rate measured during the transient, depending on if P1 binding is regulated or not.

**Table S1.** Parameters of the fitting with the model with regulation of 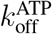 and 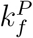. The first letter of the state conveys the configuration: nano*d*isk or vesicle. The following four letters are the receptor’s modification state.

|  |  |  |  |
| --- | --- | --- | --- |
| $k_{\text{off}}^{\text{P1}}$ | 100 s<sup>-1</sup> | $K_d^{\text{P1}}$ | 209 / 333(d) μM |
| $k_{\text{off}}^{\text{ATP}}$ | 12.8 s<sup>-1</sup> · σ | $K_d^{\text{ATP}}$ | 297 μM |
| $k_f^P$ | 21.7 s<sup>-1</sup> · σ | $G^P$ | 11.6 |

Table S1: Parameters of the fitting with the model with regulation of $k_{\text{off}}^{ATP}$ and $k_f^P$ . The first letter of the state conveys the configuration: nanodisk or vesicle. The following four letters are the receptor's modification state.
| State | [Asp] | σ |
| --- | --- | --- |
| dQEQE | 20 μM | 0.0068 |
|  | 5 μM | 0.123 |
|  | 0 μM | 0.49 |
| vEEEE | 10 μM | 0.00030 |
|  | 0 μM | 0.00177 |
| vQEQE | 100 μM | 0.0116 |
|  | 5 μM | 0.067 |
|  | 0 μM | 0.31 |
| vQQQQ | 1000 μM | 0.0176 |
|  | 0 μM | 1 |

**Figure S4:**
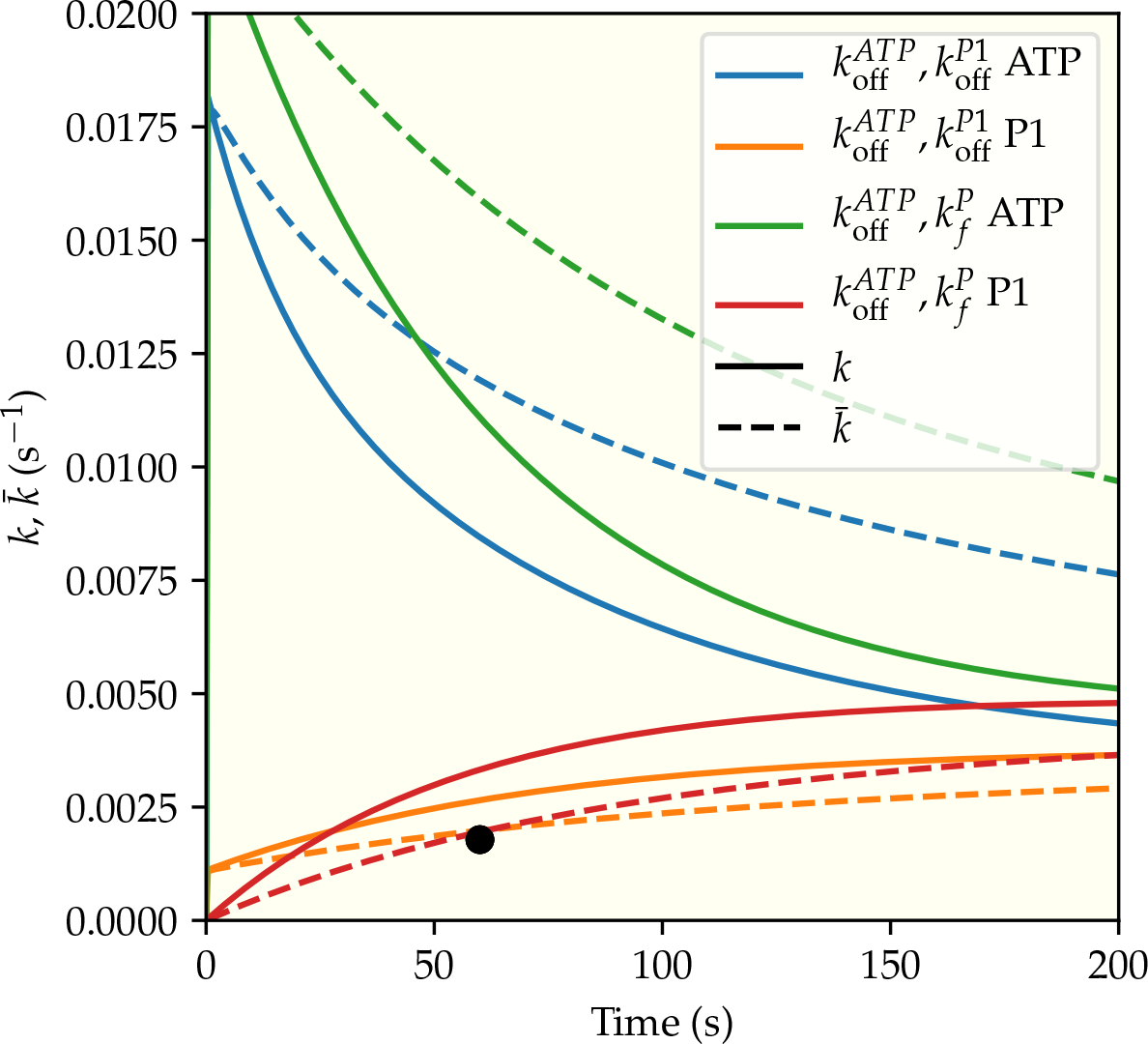
Simulations performed with the two best models in the less active of the experimental states, vesicles with EEEE receptors with [Asp]=10 μM, [ATP]=10 ^3^μM, and [P1]=25 μM the point at the lowest left corner of Fig. 3(c).

**Figure S5:**
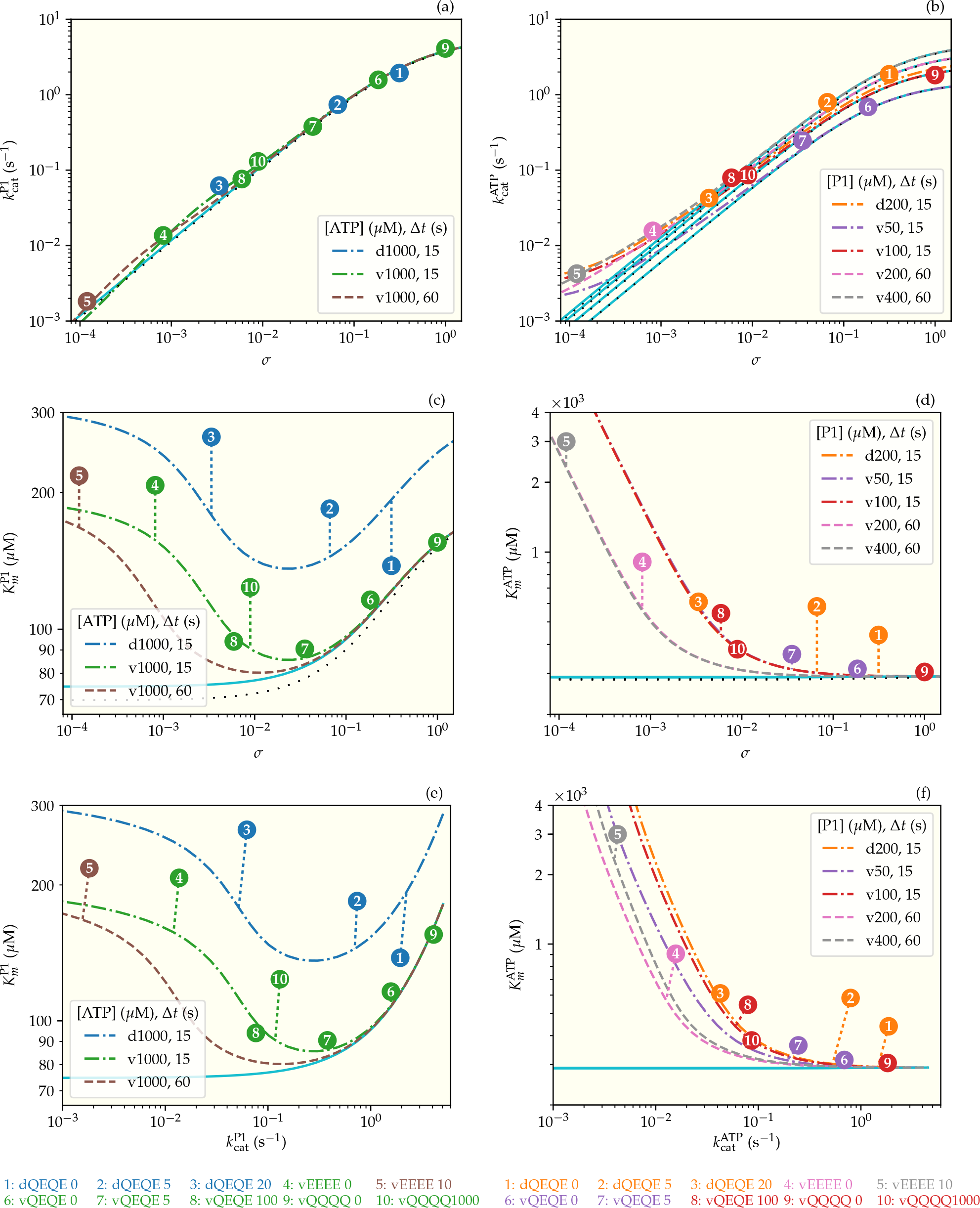
Dependence of 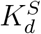 and 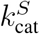 on *σ*. The curves are the parameters 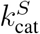 and 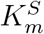 of the Michaelis-Menten equation, Eq. [1], for the phosphorylated rate predicted by the enzymatic reaction model. The parameters are calculated by fitting to curves similar to the continuous lines of Fig. 3. The simulations with Δ*t* = ∈ (steady states) are the continuous lines. Simulations for the regulatory signal *σ* ranging from 10^−3^ to 10^2^ were used to generate the curves of 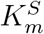 and 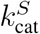 as functions of *σ* of plots (a), (b), (c), and (d).

